# Exploring Proteomes of Robust *Yarrowia lipolytica* Isolates Cultivated in Biomass Hydrolysate Reveal Key Processes Impacting Mixed Sugar Utilization, Lipid Accumulation, and Degradation

**DOI:** 10.1101/2021.04.12.439577

**Authors:** Caleb Walker, Bruce Dien, Richard J. Giannone, Patricia Slininger, Stephanie R. Thompson, Cong T. Trinh

## Abstract

*Yarrowia lipolytica* is an oleaginous yeast exhibiting robust phenotypes beneficial for industrial biotechnology. The phenotypic diversity found within the undomesticated *Y. lipolytica* clade from various origins illuminates desirable phenotypic traits not found in the conventional laboratory strain CBS7504, which include xylose utilization, lipid accumulation, and growth on undetoxified biomass hydrolysates. Currently, the related phenotypes of lipid accumulation and degradation when metabolizing non-preferred sugars (e.g., xylose) associated with biomass hydrolysates are poorly understood, making it difficult to control and engineer in *Y. lipolytica*. To fill this knowledge gap, we analyzed the genetic diversity of five undomesticated *Y. lipolytica* strains and identified singleton genes and genes exclusively shared by strains exhibiting desirable phenotypes. Strain characterizations from controlled bioreactor cultures revealed that the undomesticated strain YB420 used xylose to support cell growth and maintained high lipid levels while the conventional strain CBS7504 degraded cell biomass and lipids when xylose was the sole remaining carbon source. From proteomic analysis, we identified carbohydrate transporters, xylose metabolic enzymes and pentose phosphate pathway proteins stimulated during the xylose uptake stage for both strains. Furthermore, we distinguished proteins in lipid metabolism (e.g., lipase, NADPH generation, lipid regulators, β-oxidation) activated by YB420 (lipid maintenance phenotype) or CBS7504 (lipid degradation phenotype) when xylose was the sole remaining carbon source. Overall, the results relate genetic diversity of undomesticated *Y. lipolytica* strains to complex phenotypes of superior growth, sugar utilization, lipid accumulation and degradation in biomass hydrolysates.

**IMPORTANCE:** *Yarrowia lipolytica* is an important industrial oleaginous yeast due to its robust phenotypes for effective conversion of inhibitory lignocellulosic biomass hydrolysates into neutral lipids. While lipid accumulation has been well characterized in this organism, its interconnected lipid degradation phenotype is poorly understood during fermentation of biomass hydrolysates. Our investigation into the genetic diversity of undomesticated *Y. lipolytica* strains, coupled with detailed strain characterization and proteomic analysis, revealed metabolic processes and regulatory elements conferring desirable phenotypes for growth, sugar utilization, and lipid accumulation in undetoxified biomass hydrolysates by these natural variants. This study provides a better understanding of the robust metabolism of *Y. lipolytica* and suggests potential metabolic engineering strategies to enhance its performance.

## INTRODUCTION

*Yarrowia lipolytica* is an important oleaginous yeast for industrial biotechnology. Wildtype strains can accumulate a remarkable 40% of cell weight in neutral lipids from lignocellulosic biomass or agricultural wastes (1). These microbial lipids are a promising alternative to petroleum and animal oils for the sustainable production of advanced fuels and oleochemicals. In addition, *Y. lipolytica* is exceptionally robust to chemical inhibitors and stressful environments, which are critical biocatalyst properties to achieve sustainable production of chemicals from low-cost biomass feedstocks. It can tolerate broad pH ranges (2), high salt concentrations (3), and organic solvents (e.g., ionic liquids) (4, 5); in fact, most *Y. lipolytica* isolates exhibit robust growth in up to 60% (v/v) undetoxified dilute acid-pretreated switchgrass hydrolysates that are normally inhibitory to microbes (6). Thus, a better understanding of the mechanisms that underpin *Y. lipolytica*’ s natural robustness would not only enable development of niche strains for novel biocatalysis but would also provide fundamental knowledge that may be applied to other industrially-relevant organisms.

Recently, significant research has focused on manipulating the metabolism of the conventional laboratory strain CBS7504 (W29), isolated from a Paris sewer and well domesticated in laboratory (7), for enhanced lipid production and utilization of pentose (e.g., xylose) and hexose (e.g., glucose) sugars in inhibitory lignocellulosic biomass hydrolysates. To increase lipid production in *Y. lipolytica*, numerous metabolic engineering strategies have been implemented that successfully redirected carbon flux to lipid metabolism (8) including overexpression of lipid biosynthesis enzymes (9) and/or disruption of the competitive β-oxidation pathway (10) and altered expression of regulators (e.g., SNF1) of lipid accumulation (11). The most lipogenic *Y. lipolytica* strain reported to date achieved 90% lipid content by simultaneous restoration of leucine and uracil biosynthesis, overexpression of diacylglycerol transferase (DGA1), deletion of peroxisome biogenesis enzyme peroxin-10 (PEX10), deletion of multifunctional β-oxidation enzyme (MFE1), and optimization of culture conditions (12).

To maximize lipogenesis from biomass hydrolysates requires efficient utilization of both hexose and pentose sugars. *Y. lipolytica* does not efficiently use xylose as a sole carbon source, albeit it processes genes for the complete xylose catabolic pathway (13). Activation of this cryptic pathway has been accomplished through adaptive evolutionary approaches resulting in improved xylose utilization (14). Furthermore, overexpression of endogenous xylose catabolic genes (15-17) and heterologous expression of xylose reductase and xylitol dehydrogenase from *Scheffersomyces stipitis* (17, 18) have successfully increased xylose consumption rates. Several transporters have also been identified in *Y. lipolytica* showing increased expression levels during xylose assimilation and combinatorial overexpression of the endogenous xylitol dehydrogenase with several of these transporters has also achieved improved growth on xylose (19). While the production phenotypes are well characterized, fundamental understanding of complex phenotypes responsible for superior growth, sugar utilization, and lipid accumulation ─ or degradation ─ during fermentation of biomass hydrolysates are still lacking.

Complementary to these engineering efforts, recent investigation into the genetic diversity of undomesticated *Y. lipolytica* strains revealed emergent robust phenotypes not present in the conventional strain CBS7504. Characterization of fifty-seven undomesticated *Y. lipolytica* isolates on inhibitory undetoxified biomass hydrolysates revealed select strains with enhanced growth, lipid production, and pentose-sugar assimilation relative to CBS7504 (6). In this study, *Y. lipolytica*’ s natural genetic diversity is further explored using a combination of detailed strain characterization and proteomic analysis. Coupled together, these analyses uncover the underlying mechanisms behind these poorly understood complex phenotypes during fermentation of biomass hydrolysates. The results presented here will aid engineering efforts to better control lipid accumulation or degradation phenotypes for optimally producing advanced biofuels and/or oleochemicals from biomass hydrolysates.

## RESULTS

### Comparative genomics reveals unique genotypes of undomesticated *Yarrowia* strains

#### Phylogenetic tree of Y. lipolytica isolates shows close similarity between genomes

Phylogenetic species analysis distinguished the first evolutionary split dividing the *Yarrowia* clade into two ancestral roots (Figure 1A). The first root contained the undomesticated YB419 and the conventional strains CBS7504 and CLIB122 (a species crossed between CBS7504 and CBS6124-2 (7)). The second root contained the remaining four non-conventional isolates, depicting YB392 and YB420 as the most divergent from YB567 followed by YB566. This result was surprising since YB392, YB419 and YB420 were all isolated from corn milling plants within Illinois (20). Interestingly, the closest related species to the *Yarrowia* clade is *Sugiyamaella lignohabitans*, an efficient pentose utilizing and facultative anaerobic yeast (21).

**Figure 1.**
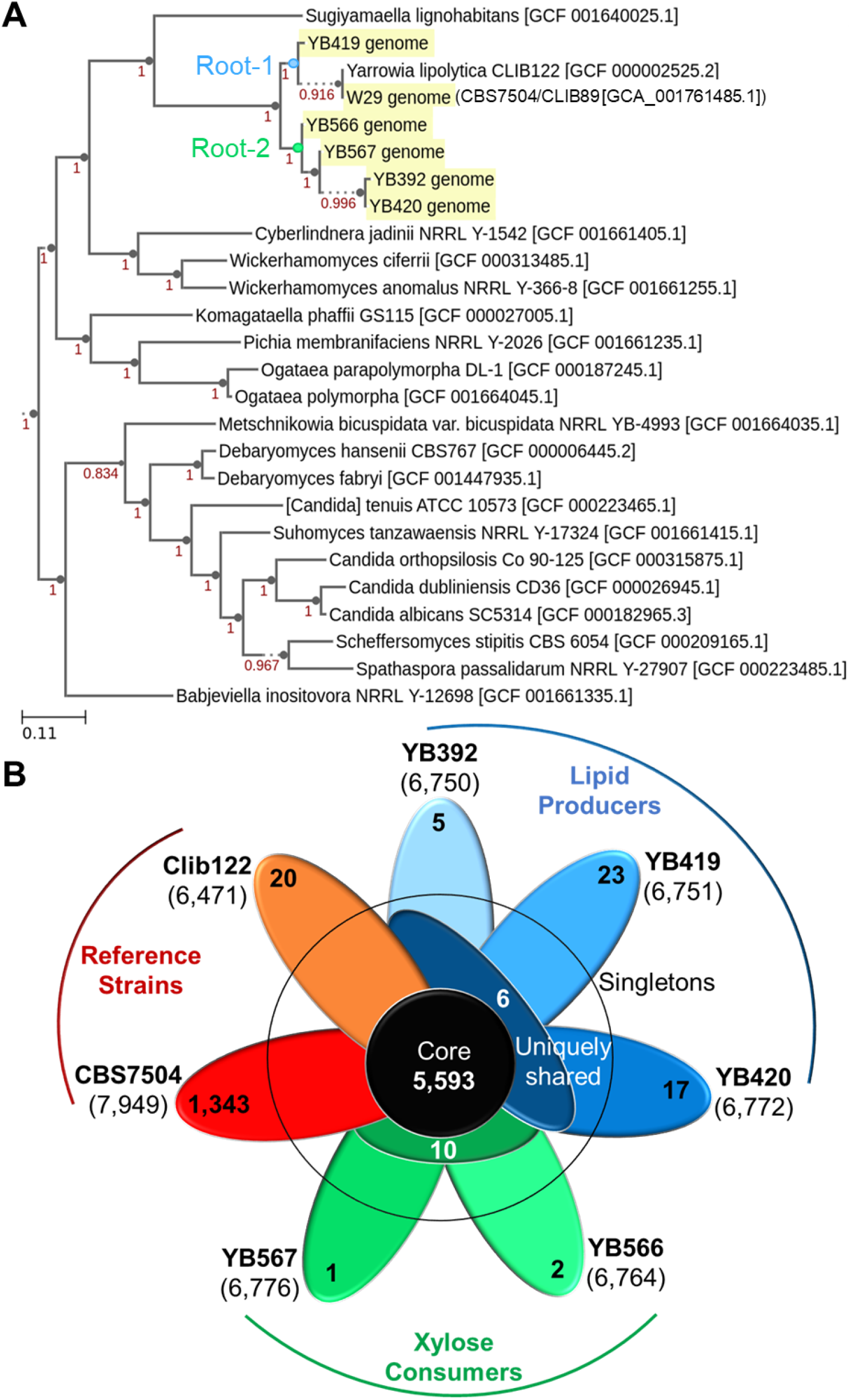
**(A)** Phylogenetic tree of *Y. lipolytica* isolates with 20 closest neighbor species. (**B**) Pangenome of *Y. lipolytica* reference strains CLIB122122 and CBS7504 and undomesticated strains YB392, YB419, YB420, YB566 and YB567. Bold, strain; parenthesis, total genes; outer petals, singleton genes, middle petals, uniquely shared genes between lipid producers (blue) and xylose consumers (green); middle circle, core genes.

#### Undomesticated Y. lipolytica isolates contain unique genes not found in conventional strains

The undomesticated strains were characterized for unique genes that may contribute to their distinctive phenotypes. A singleton gene signifies a gene appearing exclusively in one of the genomes within the pangenome (i.e., conventional strains CBS7504 and CLIB122; undomesticated strains YB392, YB419, YB420, YB566 and YB567). Of the undomesticated *Y. lipolytica* strains, YB419 contained the most singletons (23 genes) followed by YB420 (17 genes), while the remaining 3 isolates contained 5 or fewer singletons (Figure 1B, Table S1). Two of the undomesticated strains YB566 and YB567, exhibiting better xylose assimilation from switchgrass hydrolysates (SGH) (6), exclusively share 10 genes not found in other three strains. (Figure 1B, Table S2). Likewise, six genes are exclusively shared among the undomesticated strains YB392, YB419 and YB420, exhibiting better lipid accumulation (Figure 1B, Table S3) (6).

#### *Y. lipolytica* strains of different origins thrive in undetoxified biomass hydrolysates and exhibit distinct phenotypes

To better understand how the genetic diversity influences the robust phenotypes in cell growth, mixed sugar co-utilization, and lipid accumulation, we characterized CBS7504 (Figure S1), YB392 (Figure S2), YB419 (Figure S3), YB420 (Figure S4), YB566 (Figure S5) and YB567 (Figure S6) in 50% (v/v) undetoxified biomass hydrolysates. Unlike the previous studies (6), we characterized these strains in computer-controlled bioreactors (Figure 2A). In general, all strains grew well in undetoxified biomass hydrolysates. However, there were distinct phenotypes among strains associated with cell growth, mixed sugar co-utilization, and lipid accumulation. Here, two representative *Y. lipolytica* strains, CBS7504 and YB420, were selected for in-depth analysis due to their distinctive differences in xylose and lipid metabolism and members of two different ancestral roots.

**Figure 2.**
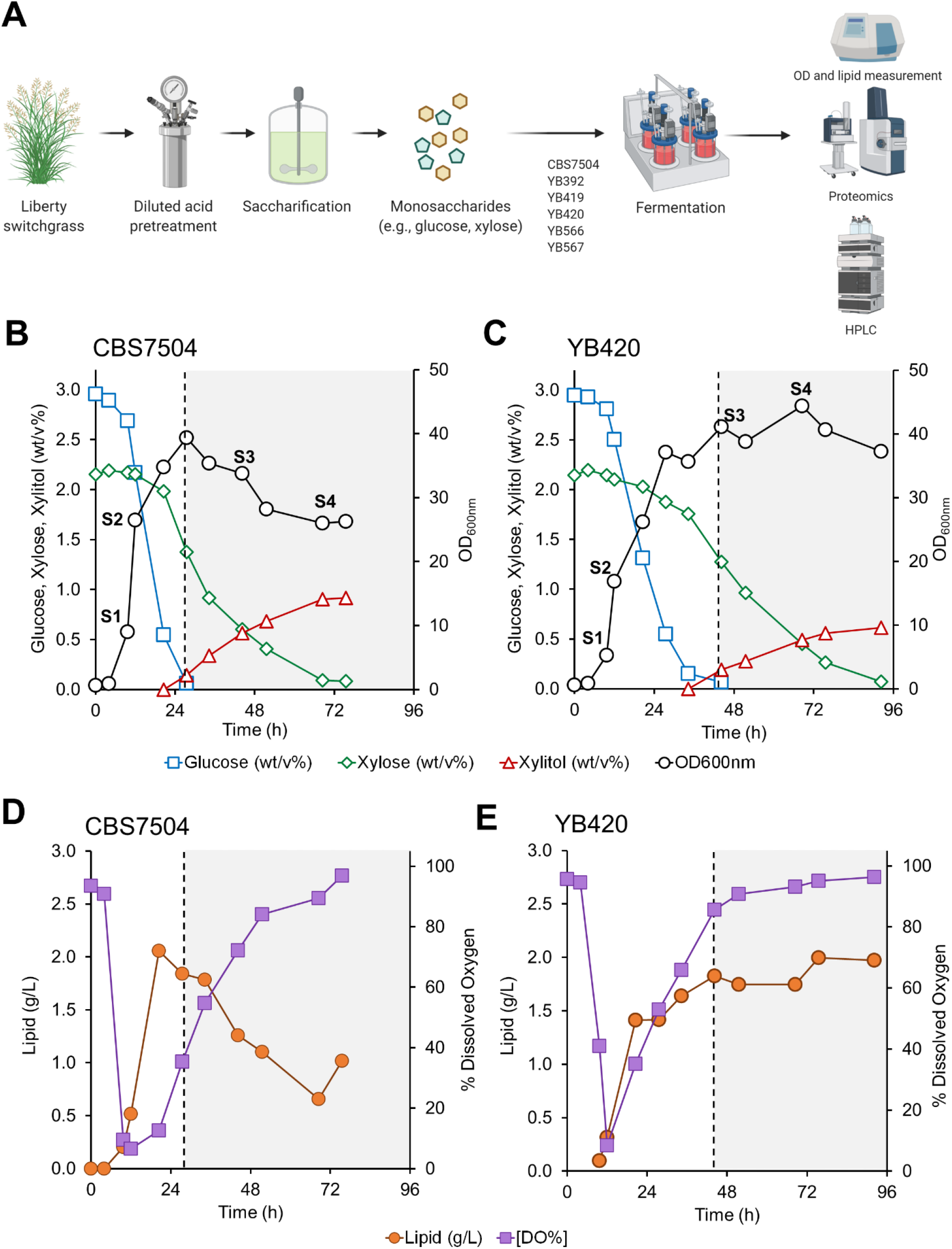
**(A)** Workflow of bioreactor characterization of *Y. lipolytica* isolates CBS7504 (**B, D**) and YB420 (**C, E**) in 50% SGH. Dotted lines, time of glucose depletion where xylose is the sole remaining carbon source.

The conventional *Y. lipolytica* strain CBS7504 grew well in undetoxified biomass hydrolysates, achieving maximum cell mass (39.3 ± 3.3 OD600nm) within 28 hours of fermentation from the co-utilization of 2.90 ± 0.05 (%w/v) glucose and 0.80 ± 0.03 (%w/v) xylose (Figure 2B). Lipids accumulated during growth, reaching a maximum of 2.10 ± 0.60 g/L at 21 hours into the fermentation (Figure 2D). Upon glucose exhaustion, cell mass and accumulated lipid levels steadily declined for the remaining 48 hours of fermentation despite the continued consumption of xylose. At 72 hours of fermentation, CBS7504 consumed a total of 2.10 ± 0.02 (%w/v) xylose and produced 0.91 ± 0.01 (%w/v) xylitol (yielding 0.44 ± 0.00 xylitol/xylose). The undomesticated *Y. lipolytica* strain YB420 also grew robustly in undetoxified biomass hydrolysates but showed contrasting phenotypes with CBS7504. Over 45 hours of fermentation, YB420 showed less co-utilization of glucose (2.80 ± 0.03 [%w/v]) and xylose (0.40 ± 0.03 [%w/v]) and less lipid production (1.6 ± 0.5 g/L) than CBS7504 (Figure 2C and E). However, upon glucose depletion, YB420 maintained cell mass and lipids while consuming a total of 2.1 ± 0.01 (%w/v) xylose and producing 0.61 ± 0.04 (%w/v) xylitol (yielding 0.30 ± 0.006 xylitol/xylose). The differences observed in xylose utilization to support maintenance of lipids and cell growth prompted a systems-level comparison between the two strains CBS7504 and YB420.

### Proteomic analysis reveals key processes impacting sugar utilization and lipid degradation

#### Proteome alterations in growth stages

Proteomic samples were collected during the exponential growth phase when glucose was assimilated (S1 and S2) and stationary phase when xylose was assimilated (S3 and S4) for both strains in biomass hydrolysate cultures. Differences in protein abundances were compared for S2, S3 and S4 against S1 for each strain (Figure 3A, 3B, 3D and 3E). As expected, there were only minor proteome differences between glucose assimilation phase samples (S1 and S2) (Figure 3A, 3D). However, *Y. lipolytica* strains dramatically altered their proteomes during stationary phase samples (samples S3 and S4) when xylose was assimilated, and lipid levels were maintained by YB420 but degraded by CBS7504 (Figure 3B, 3E). For CBS7504, 673 protein abundances changed throughout the stationary phase (samples S3 and S4) (Figure 3C) while 800 protein abundances were changed in YB420 (Figure 3F). Since the largest proteome differences were found between exponential (S1 and S2) and stationary (S3 and S4) phase samples, we chose to characterize xylose assimilation and lipid degradation phenotypes using proteins with significant changes in abundance at S3 and/or S4 relative to S1.

**Figure 3.**
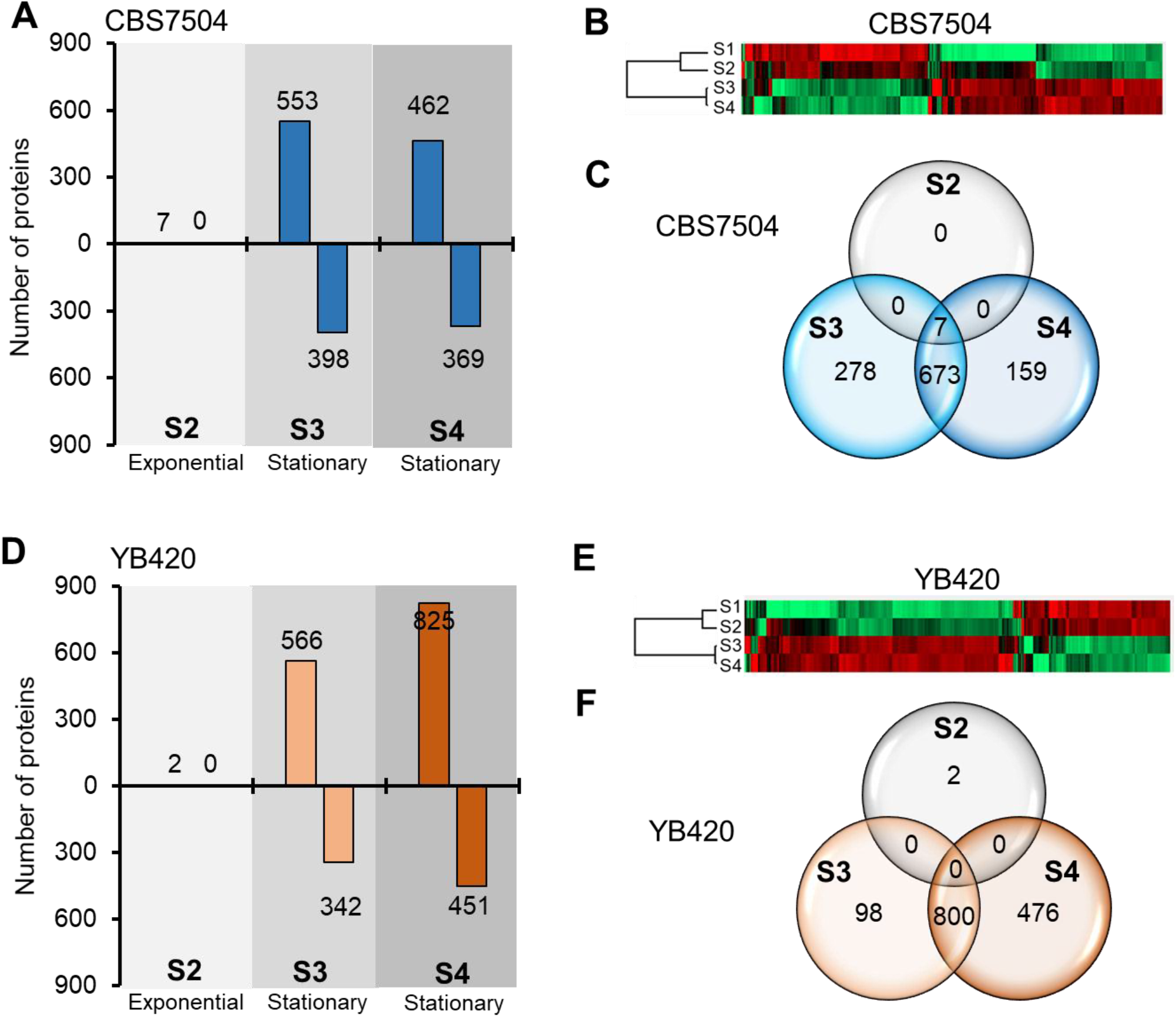
Proteomic analysis of *Y. lipolytica* (**A-C**) CBS7504 and (**D-F**) YB420 strains. (**A, D**) Number of proteins with significant abundance changes relative to S1. (**B, E**) Heatmap of significant proteins with changed abundance relative to S1. (**C, F**) Venn diagram illustrating the number of proteins with significant abundance changes between samples.

#### Proteome alterations in xylose assimilation

CBS7504 consumed xylose slightly faster than YB420 but converted more of it into xylitol (0.44 ± 0.002 %w/w of xylose) rather than maintaining cell mass or lipid content (Figure 2B, 2C). This led us to investigate protein abundances in the pentose phosphate pathway (PPP) where xylose is introduced into central metabolism. Despite the quick assimilation of xylose, during xylose assimilation (S3 and S4) CBS7504 only upregulated the protein abundance of transketolase (TKL, YALI0D02277g) in the PPP (Figure 4A, Table S4). Interestingly, CBS7504 downregulated the protein abundance of ribose-phosphate pyrophosphokinase (PRS1, YALI0B13552g) which converts ribose-5-phosphate into 5-phosphoribosyl 1-pyrophosphate (PRPP) to feed downstream biosynthetic pathways (i.e., histidine, pyrimidine and purine metabolism) associated with cell growth, correlating with the decreased cell mass, increased xylitol production and lipid degradation phenotypes of CBS7504. Meanwhile, during xylose assimilation (S3 and S4) YB420 produced less xylitol (0.30 ± 0.006 %w/x of xylose) and maintained cell mass and lipid content (Figure 2C). Regarding the PPP, six proteins were upregulated and none downregulated in YB420 during the stationary phase (Figure 4B, Table S4). Not surprisingly, these upregulated proteins include xylitol dehydrogenase (Xyl2, YALI0E12463g) and xylulokinase (Xyl3, YALI0F10923g) which together convert xylitol into xylulose-5P. Notably, YB420 also upregulated 2 proteins annotated (in panther database) for xylulose kinase (YALI0D15114g) and ribulokinase (YALI0E13321g) activities. Additionally, YB420 increased the protein abundance of D-arabinitol 2-dehydrogenase (ADH, YALI0F02211g).

**Figure 4.**
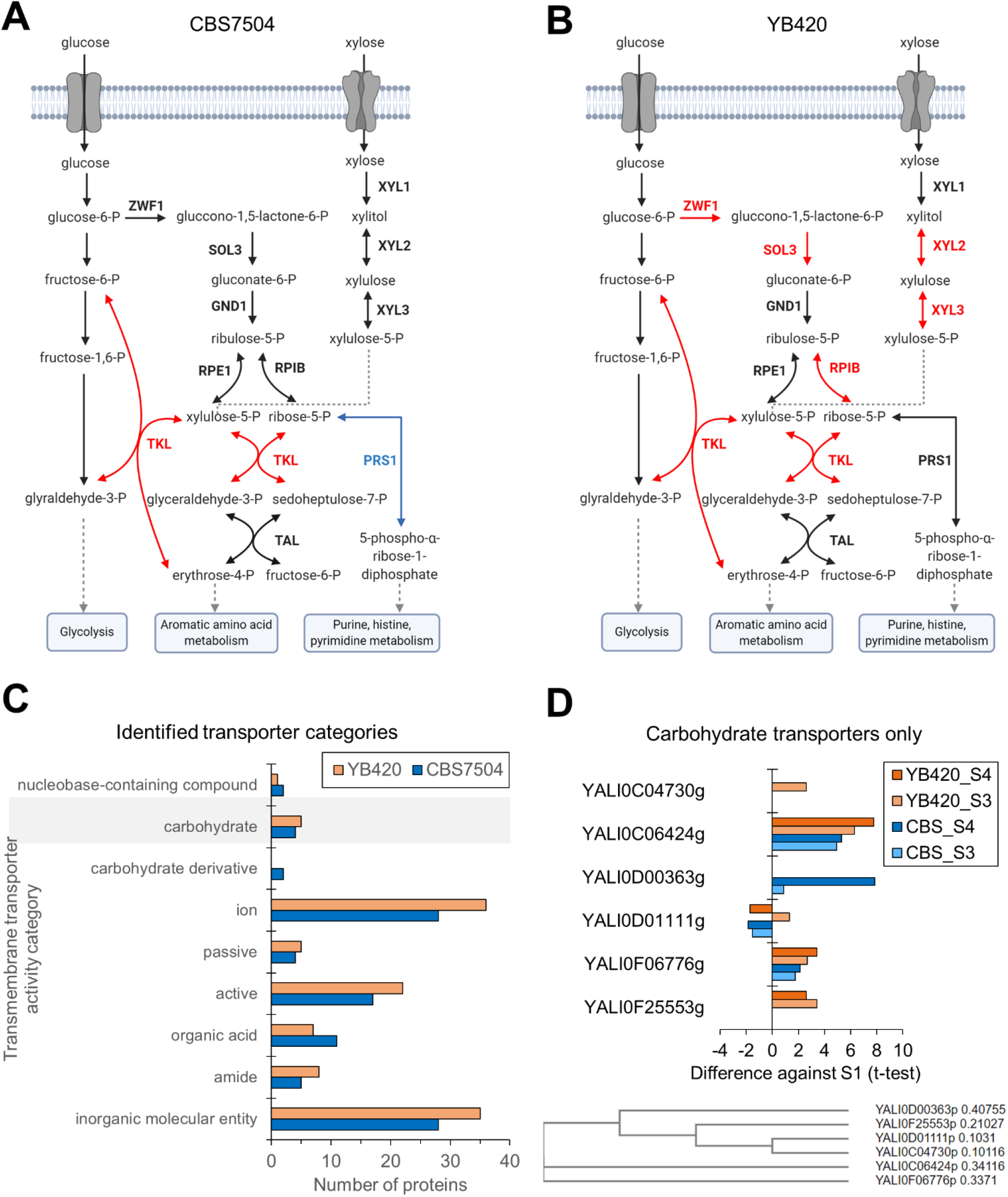
Pentose phosphate pathway proteome of (**A**) CBS7504 and (**B**) YB420. Increased protein abundance during xylose assimilation, red; decreased protein abundance during xylose assimilation, blue; pathways, rectangles; proteins, bold font; metabolites, plain text. (**C**) All transporters and (**D**) carbohydrate transporters with significant protein abundance changes in the xylose assimilation phase. CBS7504, blue; YB420, orange. Abbreviations: TKL (transketolase, YALI0D02277g); PRS1 (ribose-phosphate pyrophosphokinase, YALI0B13552g); Xyl1 (xylose reductase, YALI0D07634g); Xyl2 (xylitol dehydrogenase, YALI0E12463g); Xyl3 (xylulokinase, YALI0F10923g); ZWF1 (NADP+-dependent glucose-6-phosphate dehydrogenase, YALI0E22649g); SOL3 (6-phosphogluconolactonase, YALI0E11671g); RPIB (ribose 5-phosphate isomerase, YALI0F01628g); TAL (transaldolase, YALI0F15587g); RPE1 (ribulose-phosphate 3-epimerase, YALI0C11880g); GND1 (6-phosphogluconate dehydrogenase, YALI0B15598g).

#### Transporters

In total, 87 transporters were identified with statistically significant abundance changes when comparing stationary phase (S3 and/or S4) to exponential growth phase (S1) (Figure 4C, Table S4). While most are annotated for ion and inorganic molecular entity transmembrane transport activities, we focused on the 28 active transmembrane transporters to identify those with altered protein abundances during xylose assimilation. Specifically, six of these transporters are annotated with carbohydrate transmembrane transporter activity and have been studied for xylose assimilation in *Y. lipolytica* (Figure 4D) (14). YALI0C06424g is a carbohydrate symporter with the largest increase in abundance for both strains. This protein is similar to Snf3p and Rgt2p proteins of *Saccharomyces cerevisiae* that are involved in glucose sensing and signaling as well as fructose and mannose transport (22). Both CBS7504 and YB420 strains also increased the abundance of YALI0F06776g, an observation that is in agreement with previous transcriptomics data measuring *Y. lipolytica*’s growth response to xylose as the sole carbon source (14). Though these two transporters exhibited similar upregulation patterns during growth on xylose across both strains, albeit with varied magnitudes, other transporters were more strain specific: YALI0D00363g, YALI0F25553g, and YALI0C04730g. YALI0D00363g was strongly upregulated in S4 for CBS7504, but not in YB420, while YB420 increased the protein abundance of YALI0F25553g at both S3 and S4 and YALI0C04730g S3 but not in CBS7504. Interestingly, individual overexpression of these 3 carbohydrate symporters (YALI0D00363g, YALI0F25553g or YALI0C04730g) supported growth on plates containing xylose as the sole carbon source (19). Lastly, YALI0D01111g was downregulated in CBS7504 during xylose assimilation S3 and S4 and at S4 in YB420.

#### Proteome alterations in lipid metabolism

CBS7504 demonstrated lipid degradation while YB420 maintained lipid content during the stationary/xylose assimilation growth stage samples (S3 and/or S4) (Figures 2D, 2E, and 5A). To understand the cause of lipid degradation/maintenance, we compared proteins involved in fatty acid degradation and triacylglycerol (TAG) metabolism. In the TAG synthesis pathway, CBS7504 increased the abundance of both bifunctional glycerol-3-phosphate/glycerone-phosphate O-acyltransferase (SCT1, YALI0C00209g), converting glycerol-3-phosphate (gly-3P) into lysophosphatidic acid (LPA), and acyl-CoA dependent diacylglycerol acyltransferase I (DGA1, YALI0E32769g), which converts diacylglycerol (DAG) into TAG (Figure 5B, Table S5). Likewise, YB420 increased the of abundance DGA1, but also lysophosphatidate acyltransferase (ALE1, YALI0F19514g) and phosphatidic acid phosphohydrolase (PAP, YALI0D27016g), which together convert LPA into TAG (Figure 5C, Table S5). However, YB420 downregulated the abundance of diacylglycerol diphosphate phosphatase/phosphatidate phosphatase (LPP1, YALI0B14531g) which has the same metabolic function as PAP (KEGG E.C.3.1.3.4).

**Figure 5.**
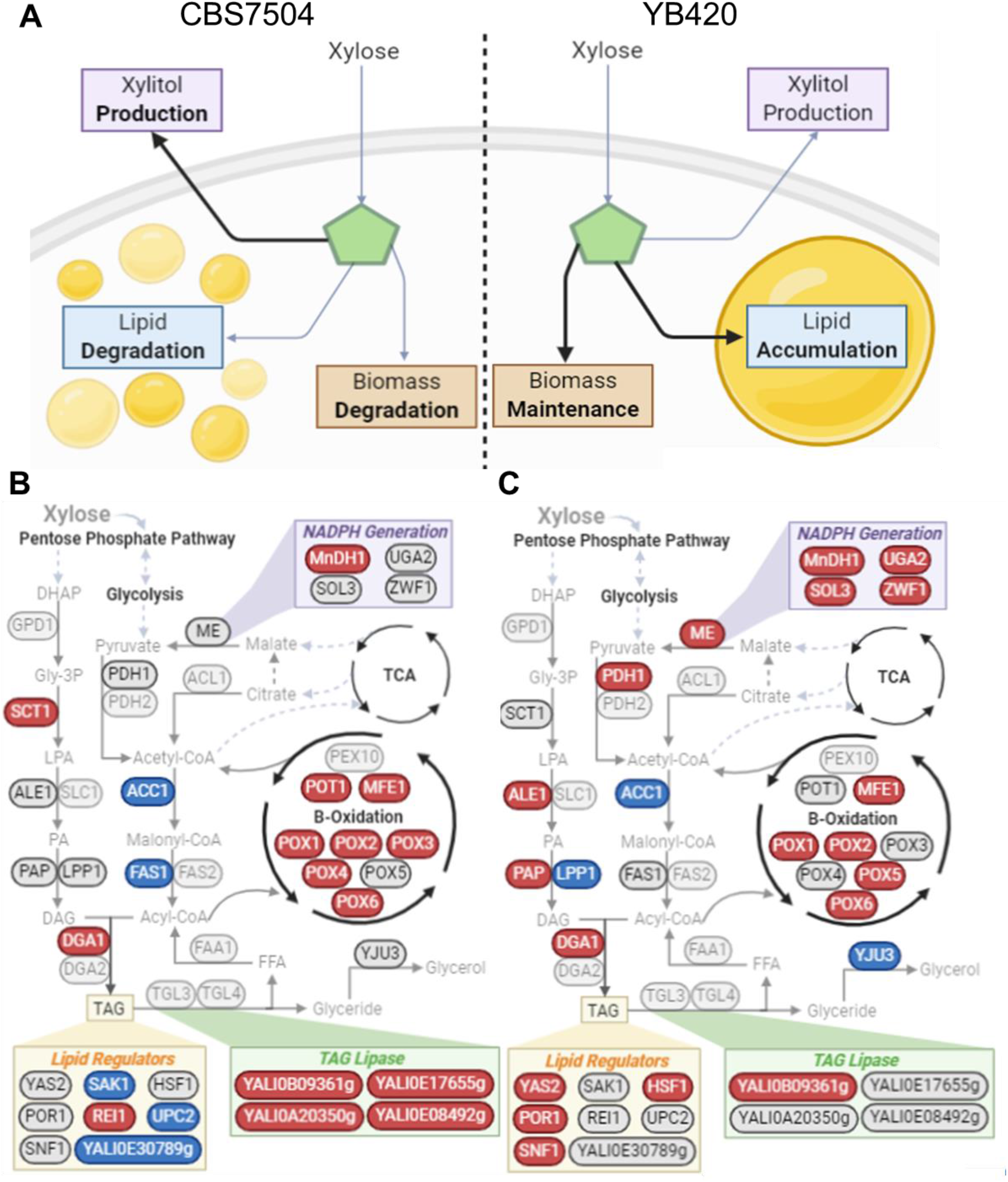
**(A)** Schematic of growth characterization phenotypes for CBS7504 and YB420 when xylose was the sole remaining carbon source. Proteomic analysis of lipid metabolism showing increased (red) and decreased (blue) protein abundance when xylose was the sole remaining carbon source for CBS7504 (**B**) and YB420 (**C**). Proteins, circles; metabolites, regular font; pathways, bold font. Abbreviations: TAG (triacylglycerol); DHAP (dihydroxyacetone phosphate); Gly-3P (glycerol-3-phosphate); LPA (lysophosphatidic acid); PA, phosphatidic acid; DAG, diacylglycerol; PL, phospholipid; TAG, triacylglycerol; FFA, free fatty acid; TCA, tricarboxylic acid cycle; GPD1 (glycerol-3-phosphate dehydrogenase, YALI0B02948g); SCT1 (bifunctional glycerol-3-phosphate/glycerone-phosphate O-acyltransferase, YALI0C00209g); ALE1 (lysophosphatidate acyltransferase, YALI0F19514g); SLC1 (1-acyl-sn-glycerol-3-phosphate acyltransferase, YALI0E18964g); PAP (phosphatidic acid phosphohydrolase, YALI0D27016g); LPP1 (diacylglycerol diphosphate phosphatase/phosphatidate phosphatase, YALI0B14531g); DGA1 (acyl-CoA dependent diacylglycerol acyltransferase I, YALI0E32769g); DGA2 (acyl-CoA dependent diacylglycerol acyltransferase II, YALI0D07986g); TGL3 (triacylglycerol lipases 3, YALI0D17534g); TGL4 (triacylglycerol lipase 4, YALI0F10010g); YJU3 (monoglycerol lipase, YALI0C14520g); FAA1 (long-chain acyl-CoA synthetase, YALI0D17864g); FAS1 (fatty acid synthase 1,YALI0B15059g); FAS2 (fatty acid synthase 2; YALI0B19382g); ACC1 (acetyl-CoA carboxylase, YALI0C11407g); PDH1, PDH2; ACL1 (ATP-citrate lyase, YALI0E34793g); ME (malic enzyme, YALI0E18634g); MnDH1 (Mannitol dehydrogenase, YALI0B16192g); UGA2 (Succinate semialdehyde dehydrogenase, YALI0F26191g); SOL3 (6-Phosphogluconolactonase, YALI0E11671g); ZWF1 (NADP+-dependent glucose-6-phosphate dehydrogenase, YALI0E22649g); POT1 (3-oxyacyl-thiolase, YALI0E18568g); MFE1 (multifunctional β-oxidation protein, YALI0E15378g); PEX10 (peroxin-10, YALI0C01023g); POX1 (acyl-coenzyme A oxidase 1, YALI0E32835g); POX2 (acyl-coenzyme A oxidase 2, YALI0F10857g); POX3 (acyl-coenzyme A oxidase 3, YALI0D24750g); POX4 (acyl-coenzyme A oxidase 4, YALI0E27654g); POX5 (acyl-coenzyme A oxidase 5, YALI0C23859g); POX6 (acyl-coenzyme A oxidase 6, YALI0E06567g); YAS2 (HLH transcription factor, YALI0E32417g); POR1 (YALI0D12628g); SNF1 (YALI0D02101g); SAK1 (YALI0D08822g); REI1 (YALI0B08734g); HSF1 (YALI0E13948g); UPC2 (YALI0B15818g);

Considering the β-oxidation pathway, both strains showed increased protein abundance of acyl-coenzyme A oxidases 1 (POX1, YALI0E32835g), 2 (POX2, YALI0F10857g), and 6 (POX6, YALI0E06567g) and multifunctional β-oxidation protein (MFE1, YALI0E15378g) (Figure 5A, 5B). However, CBS7504 also showed increased abundance of POX3 (YALI0D24750g), POX4 (YALI0E27654g) and 3-oxyacyl-thiolase (POT1, YALI0E18568g), all involved in the breakdown of TAGs into free fatty acids (FFA). Both strains decreased the abundance of acetyl-CoA carboxylase (ACC1, YALI0C11407g) but CBS7504 also decreased abundance of fatty acid synthase subunit 1 (FAS1, YALI0B15059g). Interestingly, YB420 increased the abundance of malic enzyme (ME, YALI0E18634g), which generates NADPH pools that are required for the FAS complex in oleaginous organisms (23), but shows little to no involvement in lipid production in *Y. lipolytica* (24-26).

#### Lipase

In total, ten lipases were identified with statistically significant abundance changes during the stationary phase (S3 and/or S4) relative to exponential growth phase (S1) (Figure 5B, 5C; Table S5). While none include the well-studied triacylglycerol lipase 3 (TGL3, YALI0D17534g) or TLG4 (YALI0F10010g), four of the identified lipases are annotated for TGL activity (PTHR23025). Interestingly, CBS7504 increased the protein abundance of all four TGLs in stationary phase while YB420 only increased the protein abundance of one.

#### NADPH generation

We identified five differentially abundant proteins involved in the generation of NADPH, the reducing equivalent required to sustain fatty acid synthesis (Figure 5B, C; Table S5). While both YB420 and CBS7504 strains increased the protein abundance of sorbitol dehydrogenase (MnDH1, YALI0B16192g) during stationary phase. Only YB420 increased abundances of the other 4 proteins including malic enzyme (ME, YALI0E18634g), succinate semialdehyde dehydrogenase (UGA2, YALI0F26191g), 6-phosphogluconolactonase (SOL3, YALI0E11671g) and NADP-dependent glucose-6-phosphate dehydrogenase (ZWF1, YALI0E22649g).

#### Regulators of lipid synthesis

Nitrogen limitation (i.e., high carbon to nitrogen ratio) is a common strategy to increase lipid synthesis from glucose (27), and numerous studies have reported regulators of lipid accumulation and genes affected by nitrogen limitation. Our analysis identified eight of these regulators with statistically significant abundance changes during the stationary phase (S3 and/or S4) relative to exponential growth phase (S1) – all of which have been previously reported to influence lipid accumulation (11, 28, 29). Of these, YB420 increased the protein abundance of HLH transcription factor YAS2 (YALI0E32417g), a subunit of the SWI/SNF chromatin remodeling complex POR1 (YALI0D12628g), AMP-activated serine/threonine protein kinase SNF1 (YALI0D02101g) and heat shock transcription factor HSF1 (YALI0E13948g) (Figure 5C, Table S5). Meanwhile, CBS7504 increased the protein abundance of cytoplasmic pre-60S factor REI1 (YALI0B08734g) and decreased the protein abundance of a zinc finger protein (YALI0E30789g), sterol regulatory element binding protein UPC2 (YALI0B15818g) and SNF1-activating kinase 1 SAK1 (YALI0D08822g) (Figure 5B, Table S5).

#### Proteome alterations in regulatory elements

In total, 46 gene-specific regulator proteins (from panther and TFDB) were identified with statistically significant abundance changes during the stationary phase (S3 and/or S4) relative to exponential growth phase (S1) (Figure 6, Table S6). Thirteen of these regulatory proteins had significant changes in both CBS7504 and YB420 (Figure 6). The protein with the largest increase in abundance, YALI0C07821g, is annotated as glucose transport transcription regulator RGT1-related (PTHR31668) but only has the conserved domain, GAL4 (smart00066) with *S. cerevisiae* RGT1p. Interestingly, four of these regulatory proteins have distinct, strain specific abundance patterns: RFX1, ASG1, STP1, and FHL1. RFX1, regulatory factor X in *S. cerevisiae*, is a major transcriptional repressor of DNA-damage-regulated genes (30). ASG1, an activator of stress-related genes, activates genes in β-oxidation, glucogenesis, glyoxylate cycle, triacylglycerol breakdown, peroxisomal transport, and helps assimilate fatty acids in *S. cerevisiae* (31). STP1, involved in species-specific tRNA processing, activates transcription of amino acid permease genes and is directly involved in pre-tRNA splicing in *S. cerevisiae* (32, 33). Finally, FHL1 (fork-head like) in *S. cerevisiae* functions as a transcription regulator of ribosomal protein transcription (34).

**Figure 6.**
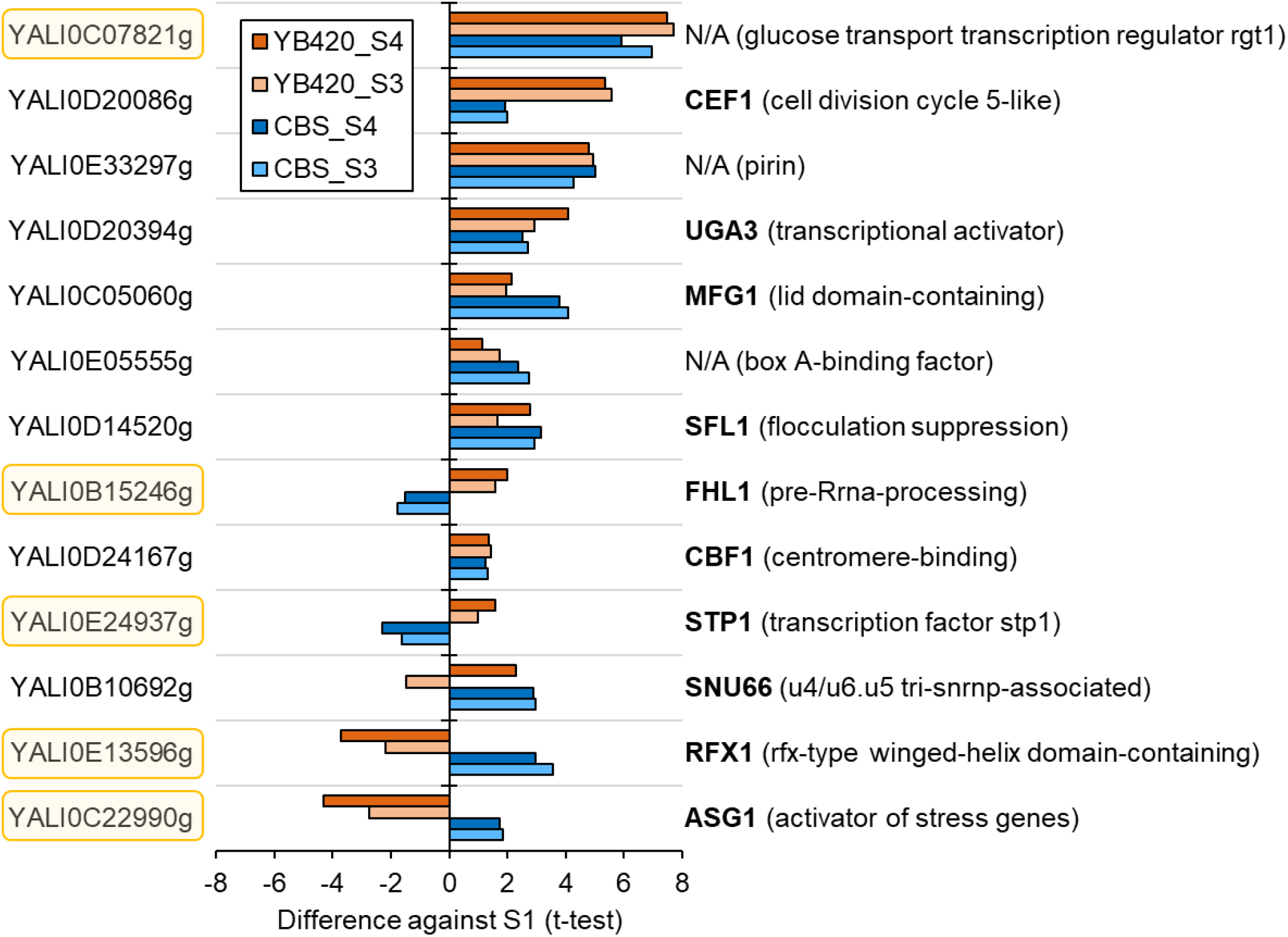
Regulator proteins with significant protein abundance changes in the xylose assimilation phase. CBS7504, blue; YB420, orange.

## DISCUSSION

Lipid accumulation and lipid degradation patterns from cultures fermenting non-preferred sugars prevalent in biomass hydrolysates (i.e., xylose) are complex phenotypes making them difficult to control and engineer in *Y. lipolytica* (15, 35, 36). By comparing proteomes of natively robust undomesticated *Y. lipolytica* strain YB420 with the conventional strain CBS7504, we identified key proteins supporting cell growth and lipid accumulation with xylose as the sole remaining carbon source.

Once all the glucose was consumed, YB420 continued to accumulate lipids and sustained cell mass from xylose, while CBS7504 degraded lipids, decreased cell mass and produced more xylitol (Figure 2). This more efficient use of xylose, demonstrated by YB420, is supported by the greater number of PPP proteins upregulated during xylose assimilation, including Xyl2 and Xyl3 which are critical for flux of xylose through the PPP (Figure 4B). Meanwhile, the almost unchanged abundances of proteins found in CBS7504 in the PPP agree with the increased xylitol secretion, decreased cell mass and lipid degradation phenotypes observed (Figure 4A). Interestingly, both strains showed similar xylose uptake profiles despite varying cell mass and lipid profiles. This suggests that transporters YALI0C06424g and YALI0F06776g are likely responsible and/or specific for xylose uptake, as indicated by increased protein abundances for both transporters across each strain only during the xylose assimilation phase (Figure 4D).

In lipid metabolism, YB420 increased proteins involved in TAG biosynthesis while CBS7504 increased proteins involved in β-oxidation and TAG lipase activity, strongly supporting the lipid maintenance and lipid degradation phenotypes of YB420 and CBS7504, respectively (Figure 5). Accordingly, YB420 increased the abundance of NADPH-generating enzymes which supply critical reducing-equivalents for lipid synthesis and xylose assimilation, including SOL3, ZWF1, ME and UGA2. Previously, overexpression of SOL3 increased lipid yield, titer and content (29). SOL3 does not produce NADPH directly, but instead catalyzes the intermediate reaction of the oxidative PPP that feeds into NADPH-producing enzymes, ZWF1 and NADP+-dependent 6-phosphogluconate dehydrogenase (29). Additionally, the promoters of ZWF1 along with ME and UGA2 exhibited increased expression levels in response to nitrogen limitation, a condition which results in increased lipid accumulation (26). Taken together, the increased protein abundance of NADPH-producing enzymes supports the lipid maintenance phenotype observed by YB420.

Several regulators previously reported to influence lipid accumulation in *Y. lipolytica* were captured by the proteomic analysis. Of these, YB420 cultures showed increased protein abundances of YAS2, POR1, SNF1 and HSF1. In a previous report, overexpression of YAS2 did not increase lipid accumulation in glucose minimal media but did significantly increase lipogenesis when acetate was the sole carbon source, suggesting indirect involvements in lipid biosynthesis in *Y. lipolytica* (29). Overexpression of POR1 in *Y. lipolytica* resulted in ∼18% increased lipid content in glycerol media but showed growth defects in glucose media (28). Conversely, deletion of SNF1 was reported to increase fatty acid accumulation without the need for nitrogen limitation (11) and overexpression of HSF1 resulted in decreased lipid accumulation with glycerol as the sole carbon source (28). We also identified one regulator (REI1) with increase protein abundance and three regulators (YALI0E30789g, SAK1 and UPC2) with decreased abundance by CBS7504 during lipid degradation. Previously, overexpression of UPC2 decreased lipid accumulation while overexpression of YALI0E30789g and REI1 increased lipid accumulation, which does not support the lipid degradation phenotype of CBS7504 in our study (28). Furthermore, deletion of SAK1 has been shown to result in increased fatty acid content (11) but in our study, decreased SAK1 protein levels were accompanied by lipid degradation in CBS7504.

While some unique genes between CBS7504 and YB420 were discovered by comparative genomics, our proteomic analysis did not identify any of them associated with the xylose or lipid metabolic phenotypes observed. This evokes the question: what underlying genotypes are causing differences in these phenotypes? While our proteomic analysis suggests regulation plays a significant role in affecting the phenotypes of the strain variants, future investigation should illuminate alterations in genome arrangement, epigenetics, and/or variants in promoter regions that could cause phenotypic divergence between CBS7504 and YB420.

In conclusion, our study highlights the regulation machinery of pentose and lipid metabolism in *Y. lipolytica* variants is complex and multifaceted, with many aspects remaining to be discovered and elucidated. Our characterization of *Y. lipolytica* isolates with phenotypic and genetic diversity, however, sheds light on those proteins supporting lipid accumulation or degradation during fermentation of non-preferred biomass sugar xylose, useful for targeted strain engineering for effective conversion of biomass hydrolysates to fuels and chemicals.

## MATERIALS AND METHODS

### Strains

*Y. lipolytica* strains CBS7504, YB392, YB419, YB420, YB566 and YB567 were used. CBS7504 is from CBS-KNAW Culture Collection in Utrecht, the Netherlands. All other strains are from ARS (NRRL) Culture Collection, Peoria, IL. The strains were stored in 20% glycerol at -80°C.

### Medium and culturing conditions

#### Switchgrass hydrolysate preparation

Liberty switchgrass, that had been pelleted and cut with a 4mm knife mill, was used. The biomass was hydrolyzed at 20% solids w/w (6). A 20-gram dry weight of biomass was added to stainless steel vessels. Then 80 mL of 0.936 % sulfuric acid w/ 3.72 g/L Pluronic F-68 was added to each vessel. Eleven vessels per oven run were filled. The 12^th^ vessel was filled with 80 mL of water and contained the thermocouple. The vessels were placed in a Mathis Labomat Infrared Oven. The following settings were used for the program: Temperature = 160°C, Heat Ramp = 2.6°C, Mix settings = 50 rpm, 60 seconds to the left and 60 seconds to the right, and cooling temperature = 40°C. Once the vessels have cooled, 4.0 mL of 1.0 M citrate buffer was added to each vessel. Then pH of the pretreated biomass was adjusted to 4.5-5.0 with 30% calcium hydroxide. The vessels were placed back in the Mathis oven for mixing at room temperature. The contents of 11 vessels were transferred to a Fernbach flask with a solid rubber stopper. The following enzymes were added: 29.7 mL Cellic Ctec3 and 5.5 mL Cellic NS-22244. The Fernbach was incubated at 50°C with shaking at 125 rpm for 3 days. After 3 days the solids were removed using a 0.2 μm filter unit. The liquid fraction was stored at 4°C. Multiple batches were made over the course of a week. The batches were pooled. The liquid was then pH adjusted to 6.0 using 10N sodium hydroxide. After pH adjustment the SGH was filter sterilized and frozen at -20°C until week of use. The SGH was thawed at 4°C overnight. Prior to use, it was amended to a whole hydrolysate with 2.31 g/L (NH_4_)_2_SO_4_, 1.81g/L Difco vitamin assay casamino acids, 0.018 g/L DL-tryptophan, and 0.072 g/L L-cysteine, and then diluted to 50% v/v with water. The diluted and amended hydrolysate is referred to 50% SGH form here on.

#### Culturing conditions

The yeast stocks were streaked on YPD Agar plates and incubated at 28°C for 24-48 hours. The plates were stored at 4°C until use. YPD media, 2 mL in a 16 mL tube, was inoculated by loop for pre-seed cultures. Pre-seed cultures were incubated at 28°C with shaking at 250 rpm for 18 hours. 0.5 mL of pre-seed culture was transferred to 10 mL 50 % SGH in 50 mL baffled flasks for seed cultures. Seed cultures were incubated 24 hours at 28°C with 250 rpm. The seed cultures were centrifuged to remove supernatant and resuspended in sterile water to A_600_ = 50. 150 mL of 50% SHG was inoculated at an A_600_ = 0.75. DasGip DasBox bioreactors were used for experimental cultures. Each strain was inoculated in triplicate. The following settings were used for the bioreactors: Beginning volume = 150 mL; Vessel = 250 mL; Temperature = 28°C; pH Set Point = 6.0; Agitation = 900 rpm; Aeration = 9.0 L/h; Base/Acid control: use 2 M HCl and 2 M NaOH for automatic dosing; Data Collection = dissolved oxygen, temperature, and pH. Cognis Clerol FBA 3107 antifoam was used to control foaming. After inoculation, 200uL of antifoam was added to each vessel. Antifoam was then added by pipet as need for duration of culture growth. A 1.2 – 1.5mL sample was taken for A600, residual sugars and lipids 3 times a day. A 1.0mL aliquot was removed from each sample for residual sugars and lipid analysis. The 1.0mL aliquot was centrifuged to remove the supernatant for residual sugar analysis. The cell pellet was washed twice with deionized water and resuspended up to 1.0mL with water. The samples were frozen at -20°C until analysis. The remaining sample was diluted for A _600_ measurement. Duplicate samples (2.0 mL) for proteomic analysis were taken once the OD reaches <4.0, 2-3 hours later, at 44-48 hours and a final sample at 68-72 hours. Samples were kept cold while processing. The samples were centrifuged to remove supernatant, washed with 1.0mL of chilled water and then centrifuged again to remove water. The washed cell pellets were the stored at -80°C.

### Analytical methods

#### Lipid quantification

Lipid analysis was done using a sulfo-phospho-vanillin colorimetric assay as previously reported by Dien *et al*. (37) For each sample, 1.0mL of sulfuric acid was added to a glass tube and 50 μL of sample (diluted with water if needed). The tube was heated at 100°C in a dry bath for 10 minutes. After heating, the tubes were cooled in a room temp water bath for 10 minutes. Once cooled, 2.5 mL of the vanillin-phosphoric acid solution was added to each tube. The tubes were mixed and placed in a 37°C incubator for 15 minutes. They are then cooled in a room temperature water bath. The absorbance is then measured at 530 nm. The Vanillin-phosphoric acid solution (0.12g vanillin, 20 mL water, and 80 mL 85% o-phosphoric acid) is made fresh daily for assays. A blank with 50uL water and four calibration standards are used for the standard curve. The calibration standards are dilutions of corn oil dissolved in 2:1 (v/v) chloroform/methanol and 50 μL of each standard was processed in duplicate along with the samples.

#### Metabolites and sugar quantification

Residual sugars were measured on a Thermo High-Performance Liquid Chromatography system. The system used a Biorad HPX-87H column and a refractive index detector. The column was kept at 65°C with 0.6mL/min of 5mM sulfuric acid as a mobile phase.

#### LC/MS for proteomic analysis

*Y. lipolytica* cells harvested at the time points detailed above were resuspended in 100 mM Tris-HCl, 10 mM dithiothreitol, pH 8.0 and bead beat with 0.5 mm zirconium oxide beads in a Geno/Grinder® 2010 (SPEX SamplePrep) for 5 min at high speed (1750 rpm). Samples were adjusted to 4% SDS and incubated at 95°C for 10 min. Crude lysate was then cleared via centrifugation (21,000 x g) and quantified by corrected absorbance (Scopes) at 205 nm (NanoDrop OneC; Thermo Fisher). Samples were then treated with 30 mM iodoacetamide for 20 min at room temperature in the dark. Three hundred micrograms of crude protein were then processed by protein aggregation capture (PAC) (38). Briefly, 300 μg of magnetic beads (1 micron, SpeedBead Magnetic Carboxylate; GE Healthcare UK) was suspended in each sample and protein aggregation was induced by adjusting the sample to 70% acetonitrile. Aggregated proteins were then washed with 1 mL of neat acetonitrile followed by 70% ethanol, and aggregated protein pellet digested with 1:75 (w/w) proteomics-grade trypsin (Pierce) in 100 mM Tris-HCl, pH 8.0 overnight at 37°C and again for 4 h the following day. Tryptic peptides released from the beads were then acidified to 0.5% formic acid, filtered through a 10 kDa MWCO spin filter (Vivaspin500; Sartorius), and quantified by NanoDrop OneC.

Peptide samples were analyzed by automated 1D LC-MS/MS analysis using a Vanquish UHPLC plumbed directly in-line with a Q Exactive Plus mass spectrometer (Thermo Scientific) outfitted with a trapping column coupled to an in-house pulled nanospray emitter as previously described (39). The trapping column (100 µm ID) was packed with 10 cm of 5 µm Kinetex C18 RP resin (Phenomenex) while the nanospray emitter (75 µm ID) was packed with 15 cm of 1.7 µm Kinetex C18 RP resin. For each sample, 3 µg of peptides were loaded, desalted, and separated by uHPLC with the following conditions: sample injection followed by 100% solvent A (95% H_2_O, 5% acetonitrile, 0.1% formic acid) chase from 0-30 min (load and desalt), linear gradient 0% to 30% solvent B (70% acetonitrile, 30% water, 0.1% formic acid) from 30-220 min (separation), and column re-equilibration at 100% solvent A from 220-240 min. Eluting peptides were measured and sequenced by data-dependent acquisition on the Q Exactive MS as previously described (40).

### Bioinformatics and data analysis

#### Comparative genomics

The following genome assemblies of *Y. lipolytica* strains were downloaded from NCBI as genbank files: CBS7504/CLIB89/W29 (GCA_001761485.1), CLIB122 (GCA_000002525.1), YB392 (GCA_003367865.1), YB419 (GCA_003367925.1), YB420 (GCA_003367965.1), YB566 (GCA_003367945.1) and YB567 (GCA_003367845.1). These were imported individually into a KBase narrative as genomes and combined into a genome set using the Build GenomeSet v1.0.1 application. A phylogenetic tree was constructed using Insert Set of Genomes Into Species Tree 2.1.10 application with neighbor public genome count of 20. Orthologue genes and unique genes shared between isolates were identified with the Compute Pangenome application using the genome set of the 7 isolates as the input. Finally, Pangenome Circle Plot - v1.2.0 application was used to produce a list of singleton genes for each base genome identified from the pangenome.

#### Proteomics

MS/MS spectra were searched against the *Y. lipolytica* CLIP122 proteome (UniProt; Nov18 build) appended with non-redundant proteins from strain YB-420 and common protein contaminants using the MS Amanda v.2.0 algorithm in Proteome Discoverer v.2.3 (ThermoScientific). Peptide spectrum matches (PSM) were required to be fully tryptic with 2 miscleavages; a static modification of 57.0214 Da on cysteine (carbamidomethylated) and a dynamic modification of 15.9949 Da on methionine (oxidized) residues. Peptide spectrum matches (PSM) were scored and filtered using the Percolator node in Proteome Discoverer and false-discovery rates initially controlled at < 1% at both the PSM- and peptide-levels. Peptides were then quantified by chromatographic area-under-the-curve, mapped to their respective proteins, and areas summed to estimate protein-level abundance. Proteins without 3 valid values in a minimum of 1 biological condition were removed and remaining protein abundances were log2 transformed. Missing values were imputed to simulate the mass spectrometer’s limit of detection using Perseus v1.6.10.43 (i.e., normal distribution, width of 0.3, down shift of 1.9 and mode changed to total matrix) (41). Significant differences in protein abundance were calculated via T-test for each sample (S2, S3 and S4) against control group (S1 sample) for each strain using FDR of 0.05, 250 permutations and s0 of 1.

All raw mass spectra for quantification of proteins used in this study have been deposited in the MassIVE and ProteomeXchange data repositories under accession numbers MSV000085941 (MassIVE) and PXD020854 (ProteomeXchange), with data files available at ftp://massive.ucsd.edu/ MSV000085941.

#### Bioinformatics

Pathway proteins were annotated using KEGG database (42) and from literature sources where cited. Ontology associations and orthologs for regulator proteins were identified using panther database (43).

## ACKNOWLEDGEMENT

We would like to acknowledge financial support from the DOE BER Genomic Science Program (DE-SC0019412 and FWP 3ERKP921). We would also like to thank the DOE’s Joint Genome Institute (JGI) for generating genome sequences of undomesticated *Y. lipolytica* strains through an EMSL FICUS award (#50384). The work conducted by the U.S. Department of Energy Joint Genome Institute, a DOE Office of Science User Facility, is supported by the Office of Science of the U.S. Department of Energy under Contract No. DE-AC02-05CH11231. The views, opinions, and/or findings contained in this article are those of the authors and should not be interpreted as representing the official views or policies, either expressed or implied, of the funding agencies. We also acknowledge the excellent technical support of Rebecca Splitt for the bioreactor studies.

## Supplementary Materials

**Table S1**. Singleton genes of YB392, YB419, YB420, YB566, and YB567.

**Table S2**. Uniquely shared genes between the undomesticated strains YB566 and YB567.

**Table S3**. Uniquely shared genes between the undomesticated strains YB392, YB419, and YB420.

**Table S4**. Proteomic analysis of xylose metabolism and transporters.

**Table S5**. Proteomic analysis of lipid metabolism, lipid regulators, lipase and NADPH generating proteins.

**Table S6**. Proteomic analysis of all gene specific regulators.

**Figure S1**. Characterization of CBS7504 growing in switchgrass hydrolysate. **(A)** Optical density measured at 600nm. **(B)** Concentrations of glucose, xylose, acetate, and xylitol. **(C)** Neutral lipids. **(D)** pH. **(E)** Volumes of base and acid added. **(F)** Percent of dissolved oxygen. Data were collected from triplicate bioreactor runs.

**Figure S2:** Characterization of the undomesticated strain YB392 growing in switchgrass hydrolysate. **(A)** Optical density measured at 600nm. **(B)** Concentrations of glucose, xylose, acetate, and xylitol. **(C)** Neutral lipids. **(D)** pH. **(E)** Volumes of base and acid added. **(F)** Percent of dissolved oxygen. Data were collected from triplicate bioreactor runs.

**Figure S3:** Characterization of the undomesticated strain YB419 growing in switchgrass hydrolysate. **(A)** Optical density measured at 600nm. **(B)** Concentrations of glucose, xylose, acetate, and xylitol. **(C)** Neutral lipids. **(D)** pH. **(E)** Volumes of base and acid added. **(F)** Percent of dissolved oxygen. Data were collected from triplicate bioreactor runs.

**Figure S4:** Characterization of the undomesticated strain YB420 growing in switchgrass hydrolysate. **(A)** Optical density measured at 600nm. **(B)** Concentrations of glucose, xylose, acetate, and xylitol. **(C)** Neutral lipids. **(D)** pH. **(E)** Volumes of base and acid added. **(F)** Percent of dissolved oxygen. Data were collected from triplicate bioreactor runs.

**Figure S5:** Characterization of the undomesticated strain YB566 growing in switchgrass hydrolysate. **(A)** Optical density measured at 600nm. **(B)** Concentrations of glucose, xylose, acetate, and xylitol. **(C)** Neutral lipids. **(D)** pH. **(E)** Volumes of base and acid added. **(F)** Percent of dissolved oxygen. Data were collected from triplicate bioreactor runs.

**Figure S6:** Characterization of the undomesticated strain YB567 growing in switchgrass hydrolysate. **(A)** Optical density measured at 600nm. **(B)** Concentrations of glucose, xylose, acetate, and xylitol. **(C)** Neutral lipids. **(D)** pH. **(E)** Volumes of base and acid added. **(F)** Percent of dissolved oxygen. Data were collected from triplicate bioreactor runs.

